# A Method for Analyzing AFM Force Mapping Data Obtained from Soft Tissue Cryosections

**DOI:** 10.1101/2023.11.08.566263

**Authors:** Cydney A. Wong, Nina Sara Fraticelli Guzmán, A. Thomas Read, Adam Hedberg-Buenz, Michael G. Anderson, Andrew J. Feola, Todd Sulchek, C. Ross Ethier

## Abstract

Atomic force microscopy (AFM) is a valuable tool for assessing mechanical properties of biological samples, but interpretations of measurements on whole tissues can be difficult due to the tissue’s highly heterogeneous nature. To overcome such difficulties and obtain more robust estimates of tissue mechanical properties, we describe an AFM force mapping and data analysis pipeline to characterize the mechanical properties of cryosectioned soft tissues. We assessed this approach on mouse optic nerve head and rat trabecular meshwork, cornea, and sclera. Our data show that the use of repeated measurements, outlier exclusion, and log-normal data transformation increases confidence in AFM mechanical measurements, and we propose that this methodology can be broadly applied to measuring soft tissue properties from cryosections.

## Introduction

Atomic force microscopy (AFM) is a widely used tool enabling the study of nanoscale properties of molecules, proteins, cells, and tissues. Furthermore, AFM enables measurements across a broad range of sample sizes in both air and liquid environments (Müller and Dufrêne, 2008; Maver et al., 2016). While commonly used to characterize surface topography of nanoscale biological materials, AFM can also be used to acquire force-displacement measurements and thus gain insights into the mechanical properties of biological samples (Binnig et al., 1986; Heinz and Hoh, 1999; Butt et al., 2005; Gautier et al., 2015). For example, there is an extensive literature on the use of AFM to study cell mechanics in culture (Haase and Pelling, 2015; Kirmizis, 2010; Li et al., 2017).

Although biomechanical characterization of cultured cells is valuable, it also suffers drawbacks. Cultured cells reside in an artificial environment, and thus typically lack the full scope of interactions with other cell types and surrounding extracellular matrix (ECM) proteins that are present in their native environment. Such interactions play an important role in many physiological and pathophysiological processes and thus can impact cellular and tissue biomechanical properties. For example, changes in cell stiffness in culture may not correlate with changes in tissue stiffness due to increased ECM deposition in response to a disease or treatment (Stylianou et al., 2018). Thus, it is useful to measure the mechanical properties of tissues *in situ* when investigating different disease states or effects of potential therapeutics. Unfortunately, the heterogeneous mix of cell types and matrix components present in tissue leads to major challenges in measuring mechanical properties of complex tissue samples, and it is perhaps in part due to this reason that AFM mechanical measurements of whole tissue samples are less common than are measurements of cultured cells (Alcaraz et al., 2018). Consequently, it is important to employ suitable AFM techniques to effectively capture and account for the biomechanical complexity, including the inherent spatial heterogeneity, of tissue samples.

One way to account for such spatial heterogeneity is through force-volume mapping, i.e. taking a dense raster scan of measurements across a sample region. Force-volume mapping, also referred to as force mapping, has been used to map spatial variations in Young’s modulus in a variety of tissue types, including in stiffer, mineralized tissues like bone and cartilage (Nemir and West, 2010; Sanchez-Adams et al., 2013; Stolz et al., 2009) as well as in soft connective tissues such as muscle (Bae et al., 2016; Engler et al., 2004), liver (Calò et al., 2020; Ojha et al., 2022; Shen et al., 2020), and neural tissues (Bouchonville et al., 2016; Christ et al., 2010; Elkin et al., 2007; Menal et al., 2018). Large variations in Young’s modulus across a single tissue or sample have been observed in many of these tissue types (Bouchonville et al., 2016; Calò et al., 2020; Franze et al., 2011; Kagemann et al., 2020; Ojha et al., 2022; Roy and Desai, 2014). However, much of this work used unsectioned pieces of tissue, or used very thick sections (>100 µm); in some applications, this is feasible, but when considering small features in complex tissues, it can be extremely challenging to find the appropriate measurement location. In such situations, alternative strategies are needed.

Here we consider one such strategy, namely the use of cryosections, which allow access to very small, specific tissue regions with intricate anatomy, while preserving both intracellular and extracellular biomolecular structures, including collagen, cytoskeletal fibers, and organelles (Li et al., 2008; Graham et al., 2010). While the snap freezing and sectioning required in this method may alter mechanical properties, causing differences as compared to the *in vivo* state, snap freezing allows for long-term tissue storage, more uniform and thin sectioning, and has been widely used in the biomedical research field (Graham et al., 2010; Peña et al., 2022; Usukura et al., 2017; Wang et al., 2017). Rapid freezing and thawing has been shown to preserve biomechanical properties of tissue sections, and a consistent experimental protocol still allows researchers to compare the effects of different biological conditions or sample locations on tissue mechanical properties (Boettcher et al., 2014; Calò et al., 2020; Lopez et al., 2011; Tran et al., 2017). AFM force-displacement measurements have thus been performed on cryosections of various tissue types, including brain, heart, lens, cornea, retina, trabecular meshwork/Schlemm’s canal, and optic nerve (Franze et al., 2011; Last et al., 2010; Menal et al., 2018; Perea-Gil et al., 2015; Vahabikashi et al., 2019; Wang et al., 2018, 2017). In all these studies, individual measurements were taken in a line or in a region of interest, rather than in a raster-scan, which may not capture the spatial heterogeneity of the tissue. While a few studies have applied force mapping to tissue cryosections (Calò et al., 2020; Liu et al., 2022; Lopez et al., 2011), they have focused on high-resolution imaging and measurement of specific matrix components or cell types within the tissue. Thus, there exists a gap in the literature regarding techniques for characterizing the overall biomechanical properties of heterogeneous soft tissues that are best accessed by cryosectioning.

Here, we developed an AFM force mapping and data analysis pipeline that addresses this gap. We use this approach to characterize the biomechanical properties of cryosectioned mouse optic nerve head tissue in a repeatable and rigorous manner. We chose to test our methodology using rodent optic nerve head samples because the mouse glial lamina tissue consists mainly of astrocytes and retinal ganglion cell axons, with some blood vessels and extracellular matrix (May and Lütjen-Drecoll, 2002; Sun et al., 2009), and this diverse composition makes it a suitable model tissue to assess force mapping techniques that can be applied to other soft, heterogeneous tissues. We further test the technique on rat trabecular meshwork (TM), cornea, and sclera to show that this measurement protocol and data analysis pipeline can also be applied to other soft tissues to obtain rigorous estimations of Young’s modulus values.

## Methods

### Mouse Optic Nerve Head Samples

All animal procedures were approved by the Institutional Animal Care and Use Committee (IACUC) of the Georgia Institute of Technology, Atlanta VA Medical Center (VAMC), or University of Iowa, and were consistent with the ARVO Statement for the Use of Animals in Ophthalmic and Vision Research. Mice used in this study were bred at the University of Iowa and shipped to the Atlanta VAMC for subsequent aging and tissue preparation. Within the current scope, the genotype of the mice was presumed to be of minor relevance, but preferably reflective of a strain for which this AFM force mapping with the optic nerve might ultimately be experimentally tested. The mice utilized were from sublines of a new transgenic model in development involving manipulations to *Apbb2,* which was generated by the University of Iowa Genome Editing Facility on an inbred C57BL/6J background. Two male and four female hemizygous mice were euthanized at 10-11 months of age (<u>Supp.</u> Table 1). Mice were sacrificed via cervical dislocation.

**Table 1:** Effects of data quality filtering on measured Young’s modulus values.

|  | Values obtained using 2 measurements at each location and Cook's distance outlier removal | Values obtained from a single measurement at each location |
| --- | --- | --- |
| Geometric mean Young's modulus (kPa) | 1.51 | 1.50 |
| Multiplicative standard deviation (kPa) | 4.26 | 4.45 |
| 95% confidence interval (kPa) | [0.18, 12.89] | [0.17, 13.33] |

Eyes were carefully enucleated, embedded in optimal cutting temperature compound (OCT), snap frozen in 2-methybutane cooled with liquid nitrogen, and stored at -80°C. Sagittal 16 µm thick sections were cut on a CryoStar NX70 cryostat (Thermofisher), through the glial lamina region, placed on Superfrost Plus Gold slides (Fisher), allowed to dry and stored at -80°C. Prior to AFM measurements, the samples were thawed and the OCT washed away by submerging in PBS for at least 10 minutes at 4°C. All samples were submerged in room temperature PBS during AFM measurements.

### Optic Nerve Head Atomic Force Microscopy

Mouse optic nerve AFM measurements were performed in the glial lamina region, located within 100 µm of the posterior sclera (Sun et al., 2009) (Fig. 1A, B), and 4-11 sections were measured per eye (Supplemental Table 3). An MFD-3D AFM (Asylum Research, Santa Barbara, CA) with a 10 µm diameter spherical probe attached to a silicon nitride cantilever (0.01 N/m) was used to obtain a raster-scan of measurements (i.e., a force map) across the glial lamina. Each map covered a 40 x 40 µm area comprised of a 4 x 4 grid of points (Fig. 1C). For each measurement, the probe approach velocity was 1 µm/s, probe retract velocity was 5 µm/s, x-y velocity during force mapping was 1 µm/s, and the trigger force was 1 nN. Each force map was repeated, and, after passing quality control tests (see below), the Young’s modulus was averaged between the two measurements to estimate the stiffness at each measurement location. One eye (37146 OD) was not measured due to technical issues.

**Figure 1:**
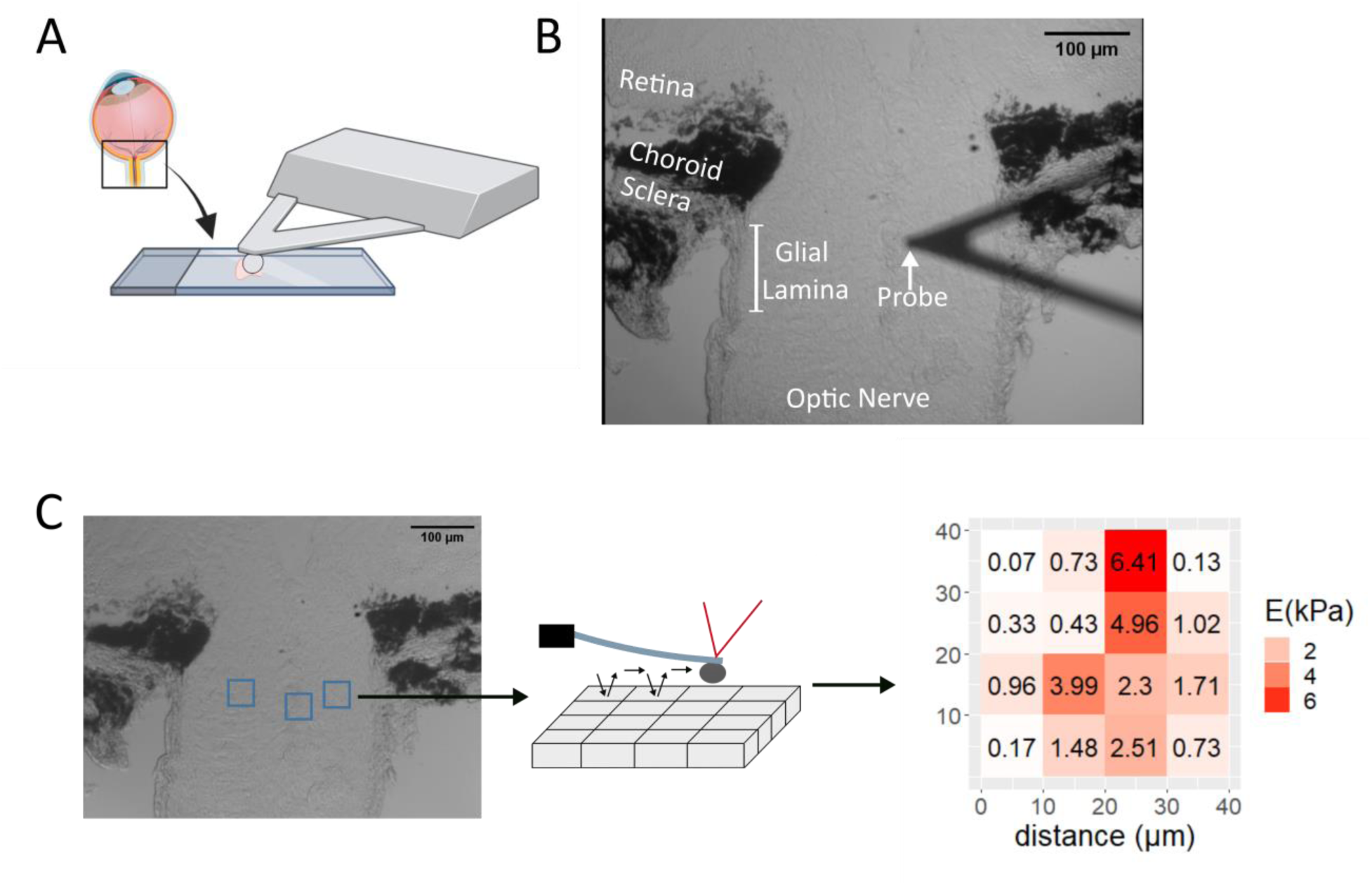
Tissue preparation and stiffness mapping methodology. **A)** After enucleation and freezing, eyes were sagittally cryosectioned as shown in the schematic, focusing on the boxed region. Sections were placed on charged slides for AFM measurement while immersed in PBS. **B)** Region of interest in a representative section as visualized by the AFM-mounted light microscope. The AFM cantilever is shown above the tissue in the glial lamina region, taken to be the region of the optic nerve within 100 µm of the posterior sclera. **C)** Overview of force mapping process. In each section. 1-3 force maps were taken in the glial lamina, each comprising a 4x4 grid of measurement, spanning a 40x40 µm area (blue boxes). An enlarged representation of the probe scanning a selected force map area is shown (middle). The resulting force curves were fit to the Hertz model and used to generate a force map. A typical map of Young’s modulus (E) values is shown (right).

### Data analysis

The Hertz model for a spherical indenter was used to fit all force-displacement curves and thereby determine the effective Young’s modulus at each location using the following formula: 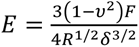 where *R* = probe radius, *δ* = indentation depth, *F* = applied force, and *υ* = Poisson’s ratio. We assumed incompressible, isotropic samples, and thus set *υ* = 0.5. Curve fitting was performed using a custom R script, and the full indentation depth was used for curve fitting, except as described below in testing.

### Outlier Removal

After fitting force-indentation curves according to the Hertz model, each curve fit was manually evaluated. Any force-indentation curves that had a sudden decrease or plateau in force during the probe approach were removed from the analysis (Fig. 2A). Additionally, if one or both force curves taken at the same location were removed due to a poor Hertzian model fit, that measurement location was entirely removed from the analysis. Furthermore, because indentation depth should be < 10% of sample thickness to avoid overestimating apparent Young’s modulus values due to substrate effects, curves with an indentation depth greater than 2 µm were removed from the analysis (Persch et al., 1994). To confirm the validity of this indentation depth cutoff value, we also took a subset of force curves from each animal, artificially truncated the force data at varying indentation depths, and calculated the fitted Young’s modulus at each indentation depth. We selected curves with an indentation depth > 2 µm and < 2 µm from the same force map within each animal for this analysis to test the indentation depth cutoff.

**Figure 2:**
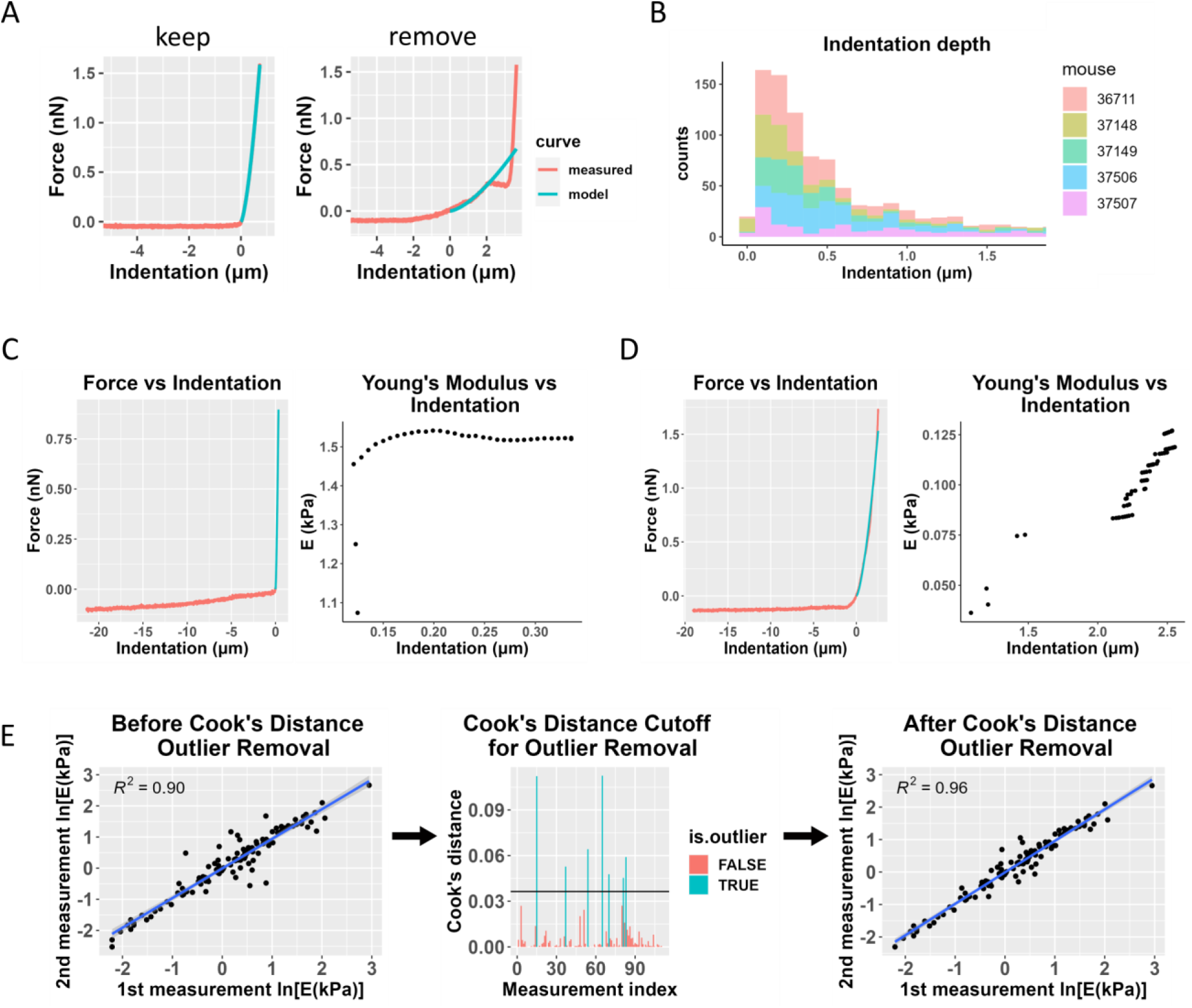
Data filtering process, including Cook’s distance for outlier removal. **A)** Representative force-indentation plots (red) illustrating curve fitting quality, with the Hertz model fit shown in blue. The “good” fit (left) demonstrates a reliable curve fit that would be retained for further analysis, while the “poor” fit (right) exhibits inadequate fitting and would be excluded from the analysis. **B)** Histogram of sample indentation depths. Each color represents data from one animal. Most measurements did not exceed a 1 µm indentation depth, and any measurements with an indentation depth greater than 2 µm were removed from the analysis. **C)** Fitted Young’s modulus values vs. indentation depth at which the force-indentation curve was truncated for analysis purposes. The plot on the left shows a sample force-indentation plot for an indentation depth < 2 µm, and the plot on the right shows the fitted Young’s modulus (E) values as a function of indentation depth for that force-indentation measurement. **D)** Similar plots are shown for a measurement from the same animal where the indentation depth exceeded 2 µm. The Young’s modulus values show much more variability and a strong dependence on the indentation depth. **E)** Overview of the use of Cook’s distance outlier removal. Log-transformed Young’s modulus estimates from the first and second measurements at the same location for one eye are plotted against each other and linearly regressed (left plot). Cook’s distance is used to determine outliers (middle plot, shown in blue), indicating discordance between repeated measurements at the same point, and the regression is re-plotted without outliers (right plot). This process was applied to data from each eye.

By repeating each force map at each measurement location, we were able to use a test-retest paradigm to verify Young’s modulus measurements. Specifically, agreement between the two measurements at the same location provided a criterion to confirm repeatability of the measurements. For each eye, the fitted Young’s modulus from the second measurement was linearly regressed on the fitted Young’s modulus from the first measurement, for all measurement locations. Cook’s distance was calculated for each data point, and measurement locations for which the Cook’s distance exceeded the cutoff *4/N*, where *N* = number of data points (Cook, 1977), were removed from the analysis (Fig. 2E). After outlier removal, two eyes (37146 OS, 37149 OS) were excluded from the analysis due to a low number of remaining measurements compared to other samples, making them unsuitable for further analysis.

### Statistics

For each mouse eye, we created histograms of Young’s modulus values after outlier removal, and a log-normal distribution was fit to Young’s modulus values using the “fitdistrplus” package in R studio (Delignette-Muller and Dutang, 2015). We confirmed the log-normal distribution with a Kolmogorov-Smirnov (K-S) test, where a critical p value of 0.05 was used. Then, we log-transformed the data and repeated the K-S test to confirm normally distributed data. We also visualized Q-Q (quantile-quantile), CDF (cumulative distribution function), and P-P (probability-probability) plots to verify that the normal distribution was a good fit to the log-transformed data.

We then pooled all 912 measured Young’s modulus values from glial lamina force mapping across nine mouse eyes and followed the same pipeline that was used for individual eyes, observing that the aggregated data also showed a log-normal distribution as judged by the K-S test, and the log-transformed aggregated data were consistent with a normal distribution by the K-S test, Q-Q plot, CDF plot, and P-P plot.

Based on the mean of the fitted normal distribution, we calculated a multiplicative (geometric) mean and multiplicative standard deviation to characterize Young’s modulus values in the non-transformed domain (Limpert and Stahel, 2011). In the same way that a normal distribution can be characterized by the arithmetic mean and standard deviation, the geometric mean and the multiplicative standard deviation, denoted by “^x^/” (i.e., times/divide) characterize the log-normal distribution.

### Rat Anterior Segment Samples

To test our data analysis pipeline in another tissue and species, we obtained rat eyes and applied a similar AFM methodology to anterior segment tissues. All animal procedures were approved by the IACUC at Emory University and the Atlanta VA Medical Center (VAMC). Eight female Brown Norway rats (Charles River), 5-6 months of age, were euthanized via inhalation of CO2 in conjunction with an approved secondary method in accordance with the Panel on Euthanasia of the American Veterinary Medical Association (AVMA) recommendations. In one eye from each animal, we followed the same freezing and embedding procedure as outlined before, and 10 µm thick sagittal cryosections of the anterior segment were placed on Superfrost Plus Gold slides (Fisher), allowed to dry, and stored at -80°C. Prior to AFM testing, the samples were submerged in PBS for at least 10 minutes at 4°C. AFM measurements were performed while samples were submerged in room temperature PBS (Fig. 3A).

**Figure 3:**
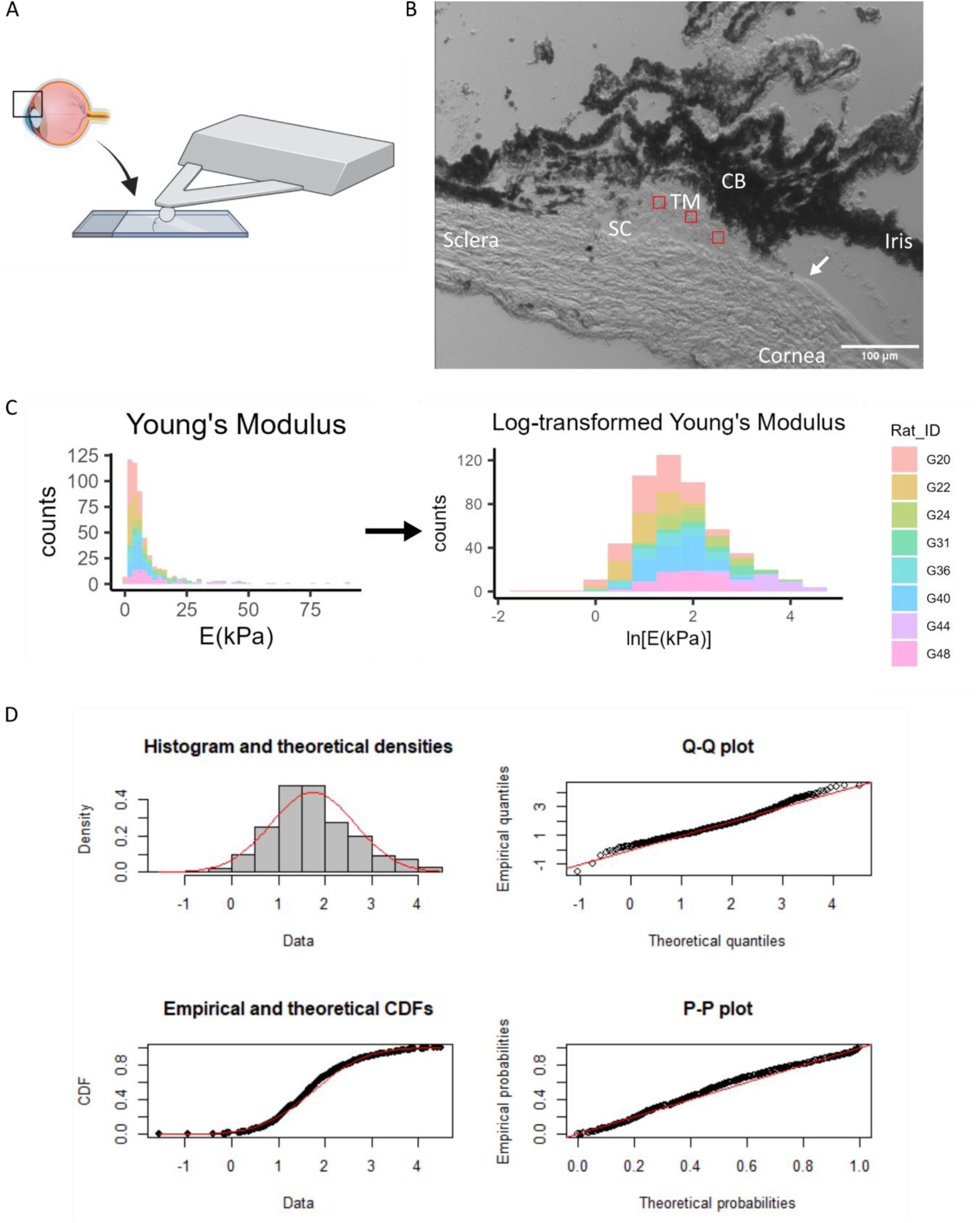
Log-transformation of Young’s modulus data from rat trabecular meshwork. **A)** Sagittal cryosections of the anterior segment were taken as shown in the schematic, focusing on the boxed region. **B)** An image of this region under the AFM probe is shown. 15 x 15 µm force maps were taken in the regions shown in red. The Schlemm’s canal and the termination of Descemet’s membrane (arrow) were the main anatomical markers used to locate the TM for force mapping. CB = Ciliary body, TM=Trabecular meshwork, SC=Schlemm’s canal. **C)** Histogram of TM Young’s modulus values from 8 rat eyes, in non-transformed and log- transformed spaces. Each color represents data from one animal. **D)** The log-transformed Young’s modulus values appeared to be well-fit by a normal distribution. Refer to Figure 3 for interpretation of graphs.

### Anterior Segment Atomic Force Microscopy

AFM force maps were acquired from the trabecular meshwork (TM), sclera, and cornea (Fig. 3B) using the same instrument and cantilever with a spherical tip as described above, except with a cantilever spring constant of 0.1 N/m. In 3-4 cryosections per eye, we took three TM force maps, each covering a 15 x 15 µm area and comprising a 4 x 4 grid of points. For the sclera and cornea, we took one force map per cryosection, covering a 20 x 20 µm area and comprising a 6 x 6 grid of points. Cornea measurements were taken on six of the eight eyes. For each measurement, the probe approach and retraction velocities were 8 µm/s, the x-y velocity during force mapping was 1 um/s, and the trigger force was 7 nN. Force indentation curves were manually inspected for goodness of fit, and curves with indentation > 1 µm (10% of section thickness) were removed from analysis. Force maps were not repeated in these tissues. We applied the same statistical methods for log-normal and normal distribution fitting as described above.

## Results

### Effect of Indentation Depth in Mouse Optic Nerve Head AFM Measurements

Across all samples, only 6.1% of data points were removed due to having an indentation depth greater than 2 µm, with most measurements remaining well below this indentation threshold (Fig. 2B). We also plotted the fitted Young’s modulus value at varying indentation depths to confirm that the reported Young’s modulus was reliable and to ensure there were minimal substrate effects at the indentation depths used in this study. The fitted Young’s modulus values were reasonably independent at indentation depths < 2 μm (Fig. 2C), and this trend was consistent for most curves in the dataset. However, we found that when the indentation depth exceeded 2 µm (10% of sample thickness), the Young’s modulus values inconsistently varied with indentation depth (Fig. 2D), justifying our decision to discard force-displacement curves with indentation depths greater than 2 μm. Simply truncating the data at 2 µm would have reduced the amount of data available for fitting, thus decreasing our confidence in the estimated Young’s modulus.

### Effects of Repeated Measurements and Cook’s Distance Outlier Removal

Approximately 16.6% of measurement pairs were removed as outliers in this study due to poor Hertz model fitting in one of the two force curves, and a further 5.2% were removed due to failing the Cook’s distance outlier criterion. Supplemental Table 3 shows the number of data points removed at each step of the pipeline. Generally, we observed good agreement between the repeated (test-retest) measurements in each force map, but removal of outliers using our Cook’s distance protocol did improve the test-retest concordance. Figure 2E shows test-retest agreement for data from a single eye, but overall, the average R^2^ values from linear regressions of test-retest Young’s modulus before and after Cook’s outlier removal were 0.82 and 0.91, respectively. However, conducting the analysis without these quality criteria did not significantly change the resulting average Young’s modulus estimate when pooling data from all the cryosections, although it did result in a slightly wider 95% confidence interval (Table 1).

### Log-normal Distribution of Mouse Optic Nerve Young’s Modulus

After log-transformation (Fig. 4A), goodness-of-fit to a normal distribution of the transformed data was evaluated by histogram, Q-Q, CDF, and P-P plots (Fig. 4B). Our analysis showed that Young’s modulus data closely followed a log-normal distribution, with overall geometric mean and standard deviation of 1.51 ^x^/ 4.26 kPa in the mouse glial lamina. The reported Young’s modulus was different when we computed the traditional arithmetic mean and standard deviation of 4.27 ± 8.06 kPa (Table 2 and Figure 4C).

**Figure 4:**
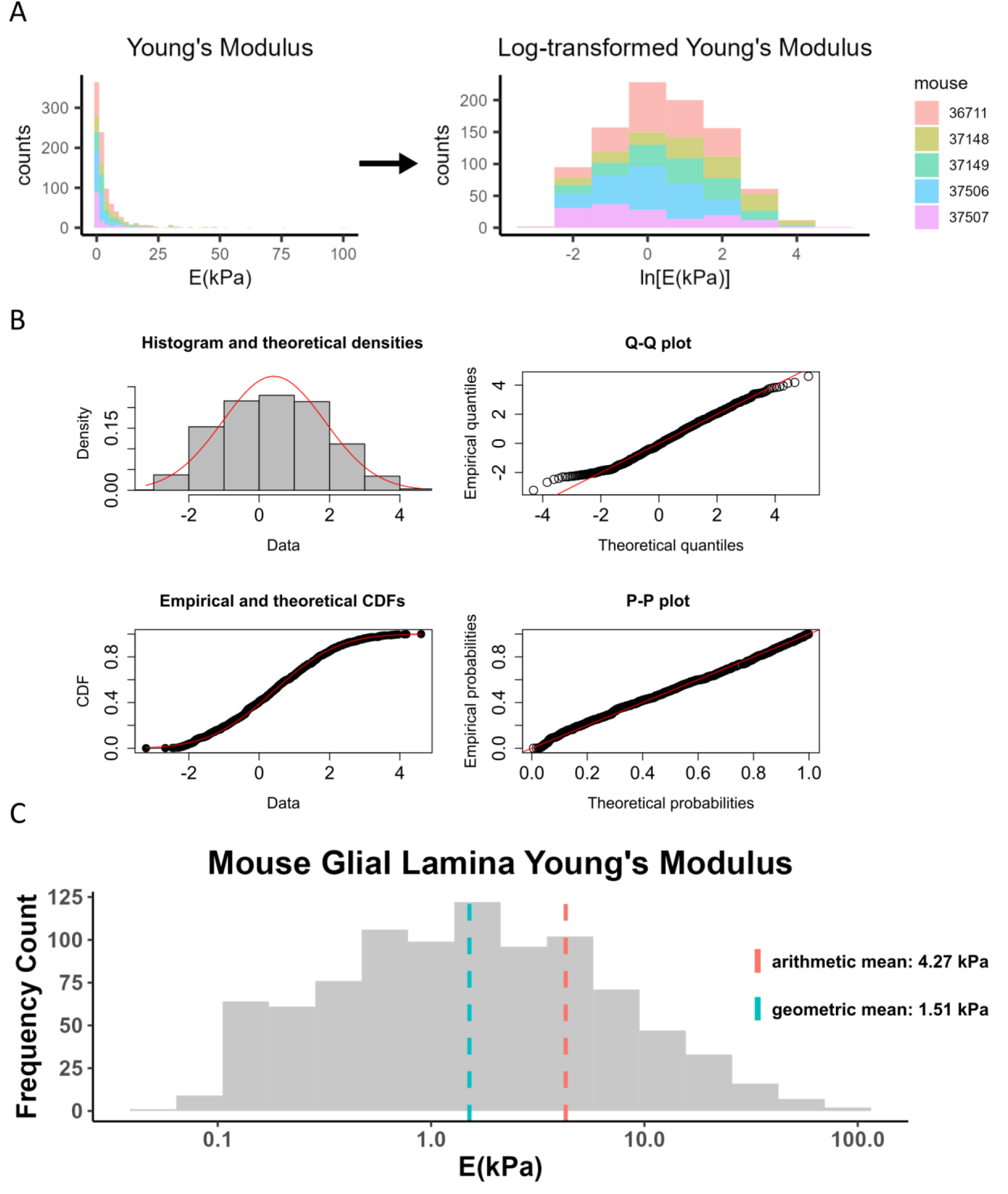
Log-transformation of Young’s modulus data from mouse glial lamina. **A)** Histogram of Young’s modulus values from 9 eyes of 5 mice. The raw data was log-transformed to obtain a distribution that appeared to be consistent with a normal distribution. Each color represents data from one animal. **B)** The log-transformed Young’s modulus values appeared to be well-fit by a normal distribution, as judged by a histogram of Young’s modulus values vs. a fitted normal distribution (top left), and by comparisons of actual and theoretical quantiles (top right), actual and theoretical cumulative distribution functions (bottom left), and actual and theoretical probability distributions (bottom right). In all four panels, actual data is in black/grey and theoretical fits are overlain in red. **C)** Histogram of Young’s modulus values showing geometric and arithmetic means. The geometric mean, indicated by the blue dashed line, better represents the data compared to the arithmetic mean. X-axis is shown on a log-scale.

**Table 2:** Young’s Modulus Summary Statistics with Log-Transformation.

|  | **Geometric mean and multiplicative standard deviation** | **Arithmetic mean ± standard deviation** |
| --- | --- | --- |
| Mean and standard deviation (kPa) | 1.51 × / 4.26 | 4.27 ± 8.06 |
| 95% confidence interval (kPa) | [0.18, 12.89] | [-11.87, 20.41] |

### Application to Other Tissues

To test whether the above data processing pipeline could be used in characterizing other soft tissues by AFM, we also measured rat TM, scleral, and corneal stiffness. A histogram of all the TM Young’s modulus values (n = 516 measurements) showed a log-normal distribution (Fig 3C), with goodness-of-fit evaluations shown in Fig. 3D. We also applied the same protocol to measure scleral and corneal stiffness in rat anterior eye cryosections and found that the scleral (Supplemental Fig. 1) and corneal (Supplemental Fig. 2) Young’s modulus values were also log-normally distributed, with the log-transformed data passing the K-S normality test. The Young’s modulus values for each region are reported in Table 3, highlighting the differences when computing the geometric and arithmetic means.

**Table 3:** Young’s Modulus Summary Statistics with Log-Transformation. Data are represented by geometric (geometric mean and multiplicative standard deviations) or arithmetic (arithmetic mean ± standard deviation) sumamry statistics.

| **Tissue** | **Calculation Method** |  |
| --- | --- | --- |
| **Rat TM** | **Geometric** | **Arithmetic** |
| Mean and standard deviation (kPa) | 5.70 × / 2.45 | 9.02 ± 11.66 |
| 95% confidence interval (kPa) | [0.94, 34.21] | [-14.30, 32.34] |
| **Rat Sclera** |  |  |
| Mean and standard deviation (kPa) | 17.19 × / 2.07 | 22.31 ± 18.55 |
| 95% confidence interval (kPa) | [4.01, 73.66] | [-14.79, 59.41] |
| **Rat Cornea** |  |  |
| Mean and standard deviation (kPa) | 6.77 × / 2.05 | 8.76 ± 6.92 |
| 95% confidence interval (kPa) | [1.61, 28.45] | [-5.08, 22.60] |

## Discussion & Conclusions

The data analysis pipeline described in this study was designed to obtain Young’s modulus values from AFM measurements of cryosectioned soft tissues in a manner that accounts for inherent tissue heterogeneity and is robust, as demonstrated through strong test-retest agreement. The proposed approach focuses on obtaining an aggregated Young’s modulus from tissue sections, rather than individual moduli from cell types or specific ECM components within a tissue. Key elements of this pipeline include careful quality control on individual force-indentation curves, the use of Cook’s distance for automated elimination of outliers, and fitting of Young’s modulus values to a log-normal distribution.

We were surprised to observe that the quality control and outlier removal aspects of this pipeline did not materially affect the overall Young’s modulus values that we estimated, with only about 20% of measurement values discarded and a modest reduction in the 95% confidence limit associated with the mean Young’s modulus value. However, this may be tissue specific, and we suggest that best practice is to apply both force curve quality control and test-retest outlier removal, at least in preliminary studies until the tissue is better characterized. This approach builds on the work of Kagemann et al., who also performed repeated force maps in human TM cryosections to show how test-retest reproducibility and Young’s modulus varied spatially (Kagemann et al., 2020). By adding the Cook’s outlier protocol with repeat force mapping as shown in Fig. 2E, we establish a consistent method for quantifying test-retest variation across an entire sample.

More significant was the use of log-normal statistics when analyzing data. AFM studies typically report Young’s modulus values as arithmetic mean ± standard deviation; however, based on the data in this study, it is clear that the geometric mean and multiplicative standard deviation better characterize the data. Indeed, the confidence intervals for Young’s modulus based on an arithmetic mean are nonsensical because they imply the existence of negative Young’s modulus values (Table 2). It was notable that Young’s modulus values from all four tissues considered in this study (mouse optic nerve head, rat TM, rat sclera, and rat cornea) followed log-normal distributions. Careful reading of the literature shows that others have reported log-normally distributed Young’s modulus values in human neuronal tissue (Bouchonville et al., 2016; Liu et al., 2022), reinforcing the suitability of this data fitting approach. More generally, the log-normal distribution commonly arises in scientific data when the measured value cannot be negative, or more generally, cannot take values below a cutoff (Limpert et al., 2001; Limpert and Stahel, 2011).

Here, we build on these studies and other existing AFM literature by measuring stiffness in tissue cryosections rather than cultured cells to capture biomechanical properties *in situ,* permitting us to link tissue stiffness to other phenotypic information, e.g., in animal models of disease. Cryosectioning of the small rodent eye allowed for precision in locating and measuring specific tissue regions, particularly critical for glial lamina measurements because different regions of the optic nerve head have different compositions.

Indentation depth is well-known to be an important parameter to consider when using the Hertz model to analyze force-displacement curves on cryosectioned tissue, since the Hertz model as used here assumes small indentation relative to the tissue thickness. Our results confirmed this requirement, showing large variations in fitted Young’s modulus values when the indentation depth was too large. While section thickness can be used as a guideline to estimate an appropriate indentation depth cutoff, the best cutoff value may be empirically determined by artificially truncating the force-displacement curve when fitting the Hertz model and observing when the fitted Young’s modulus values begin to show large variation as a function of truncated indentation depth or by applying a strain-dependent evaluation criteria (Xu et al., 2023).

Although the pipeline developed in this study used rodent ocular tissue samples, this approach should enable more consistent and repeatable AFM force measurements of soft tissue cryosections more generally.

## Acknowledgements

We thank Kelleigh Hogan for her help in handling the mice at the Atlanta VA Medical Center. This work was supported by the NIH (Diversity Supplement EY031710-01S1 [CAW], R01 EY031710-01S1 [CRE], T32 Training Grant GM145735 [CAW], R21 EY031598 [MGA], R01 EY030871 [AJF], P30EY006360 [AJF], Training Grant T32 EY007092 [NSFG]), the Georgia Research Alliance (CRE), NSF CBET 2225476 [TAS], the Alfred P. Sloan Foundation G-2019-11435 [NSFG] and the Department of Veterans Affairs Rehab R&D Service Career Development Awards to AJF (CDA-2; RX002342)

