## Supplemental Files for "A Method for Analyzing AFM Force Mapping Data Obtained from Soft Tissue Cryosections"

### Supplemental Materials

#### *Supplemental Figure 1: Young's Modulus in Rat Sclera*

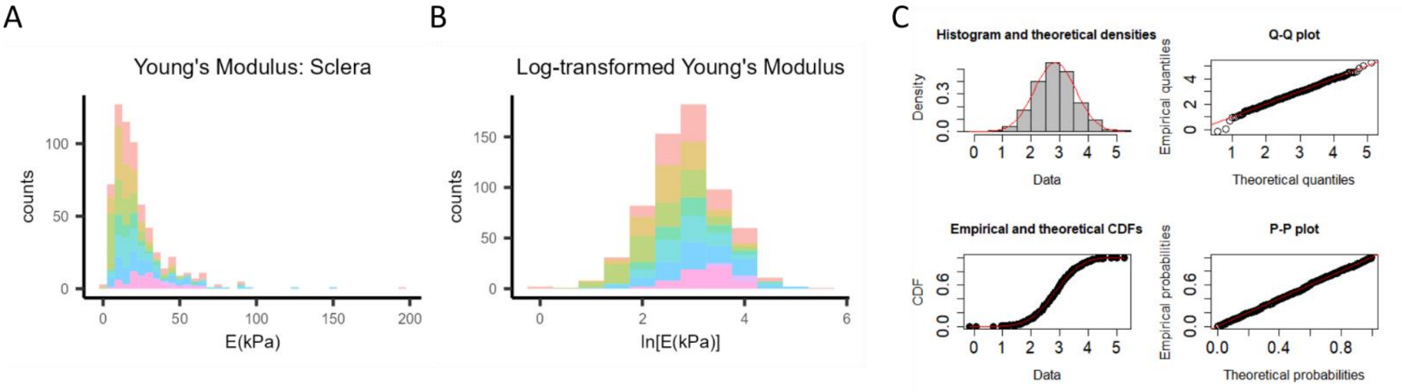


***A)*** *Histogram of scleral Young's modulus values (n=631 measurements, 8 rat eyes).* ***B)*** *Log-transformed Young's modulus data showing an approximately normal distribution.* ***C)*** *Goodness-of-fit plots comparing a normal distribution fit to the log-transformed scleral Young's modulus values shows high agreement between empirical and theoretical data. This data passed the K-S normality test. See Figure 3 for explanation of plots.*

#### *Supplemental Figure 2: Young's Modulus in Rat Cornea*

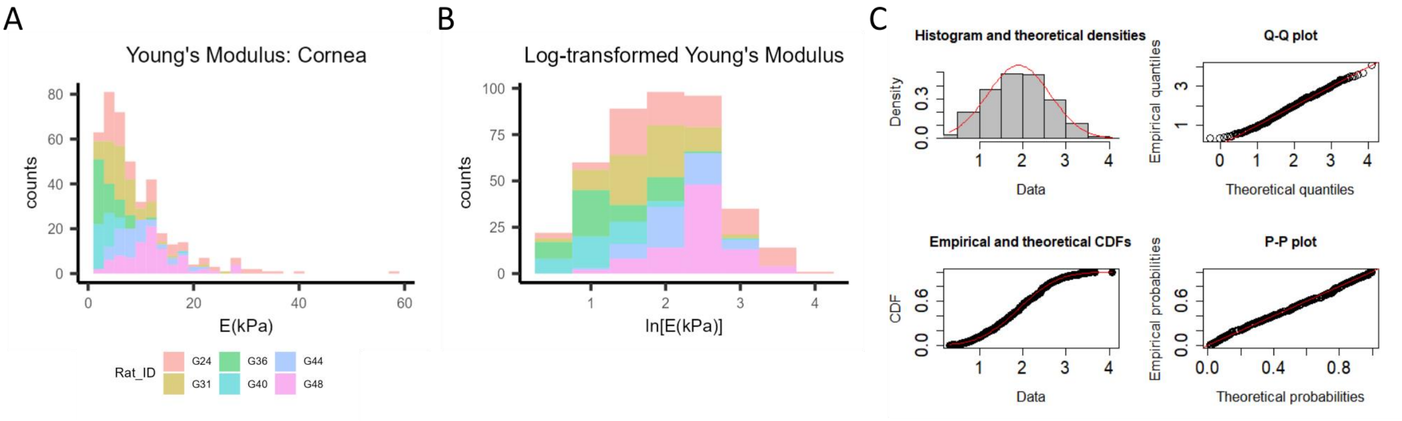


***A)*** *Histogram of cornea Young's modulus values (n=415 measurements, 6 rat eyes).* ***B)*** *Log-transformed Young's modulus data showing an approximately normal distribution.* ***C)*** *Goodness-of-fit plots comparing a normal distribution fit to the log-transformed scleral Young's modulus values shows high agreement between empirical and theoretical data. This data passed the K-S normality test. See Figure 3 for explanation of plots.*

*Supplemental Table 1: Mice used in this study*

| Mouse ID | Date of birth | Sacrifice date | Age | Sex |
|---|---|---|---|---|
| 36711 | 6/8/2021 | 5/17/2022 | 11 mo | M |
| 37146* | 9/3/2021 | 7/13/2022 | 10 mo | M |
| 37148** | 9/3/2021 | 7/13/2022 | 10 mo | F |
| 37149 | 9/3/2021 | 7/13/2022 | 10 mo | F |
| 37506 | 11/11/2021 | 9/14/2022 | 10 mo | F |
| 37507 | 11/11/2021 | 9/14/2022 | 10 mo | F |

*Both eyes removed from analysis (not enough data/technical issues)

** Only OS (left) eye used in analysis*Supplemental Table 2: Rats used in this study*

| Rat ID | Date of birth | Sacrifice date | Age | Sex |
|---|---|---|---|---|
| G20 | 01/21/22 | 08/29/2022 | 7 mo | F |
| G22 | 01/21/11 | 09/14/2022 | 7 mo | F |
| G24 | 07/03/2022 | 10/24/2022 | 3 mo | F |
| G31 | 07/03/2022 | 11/14/2022 | 4 mo | F |
| G36 | 07/03/2022 | 11/28/2022 | 5 mo | F |
| G40 | 10/03/2022 | 03/06/2023 | 5 mo | F |
| G44 | 10/03/2022 | 03/19/2023 | 5 mo | F |
| G48 | 10/03/2022 | 04/03/2023 | 6 mo | F |

*Supplemental Table 3: Optic Nerve Head Outlier Removal data*

| | | | | **Number of data points remaining after each quality control step** | | | |
|---|---|---|---|---|---|---|---|
| Mouse | Eye* | Geometric mean E (kPa) | Number of cryosections measured | Remove poor Hertz model fits | Filter for indentation < 2 µm | Filter for repeated measurements | Cook's distance outlier removal |
| 36711 | OD | 2.391 | 8 | 137 | 130 | 107 | 99 |
| | OS | 1.125 | 10 | 157 | 155 | 155 | 144 |
| 37148 | OS | 2.996 | 8 | 219 | 193 | 154 | 150 |
| 37149 | OD | 0.91 | 4 | 83 | 80 | 64 | 64 |
| | OS | 3.57 | 6 | 152 | 135 | 96 | 91 |
| 37506 | OS | 0.941 | 9 | 117 | 115 | 78 | 74 |
| | OD | 1.148 | 11 | 154 | 153 | 153 | 144 |
| 37507 | OD | 0.757 | 5 | 122 | 108 | 91 | 86 |
| | OS | 1.526 | 4 | 88 | 85 | 64 | 60 |
| **Total:** | | | 65 | 1229 | 1154 | **962** | **912** |

** OD = right eye, OS = left eye*
